# SplicingLore : a web resource for studying the regulation of cassette exons by human splicing factors

**DOI:** 10.1101/2023.06.30.547181

**Authors:** Hélène Polvèche, Jessica Valat, Nicolas Fontrodona, Audrey Lapendry, Stéphane Janczarski, Franck Mortreux, Didier Auboeuf, Cyril F. Bourgeois

## Abstract

One challenge faced by scientists from the alternative RNA splicing field is to decode the cooperative or antagonistic effects of splicing factors to understand and eventually predict splicing outcomes on a genome-wide scale. In this manuscript, we introduce SplicingLore, an open access database and web resource that help to fill this gap in a straightforward manner. The database contains a collection of RNA-seq-derived lists of alternative exons regulated by a total of 75 different splicing factors. All datasets were processed in a standardized manner, ensuring valid comparisons and correlation analyses. The user can easily retrieve a factor-specific set of differentially included exons from the database, or provide a list of exons and search which splicing factor(s) control(s) their inclusion. Our simple workflow is fast and easy to run, and it ensures a reliable calculation of correlation scores between the tested datasets. As a proof of concept, we predicted and experimentally validated a novel functional cooperation between the RNA helicases DDX17 and DDX5 and the HNRNPC protein. SplicingLore is available at https://splicinglore.ens-lyon.fr/.

## INTRODUCTION

Eukaryotic genes are transcribed into pre-messenger RNAs (pre-mRNAs) that are most of the time composed of a succession of exons separated by introns, the latter being removed by the spliceosome to form mature RNA molecules or mRNAs. The spliceosome is a large ribonucleoprotein complex which recognizes the 5’ and 3’ splice sites delimitating exons from introns and catalyzes the splicing reaction, a process that is modulated by numerous auxiliary factors, including RNA binding proteins (RBP) (1,2). The split nature of genes and the contribution of many different parameters enables alternative splicing, which is defined as the differential selection of exonic or intronic sequences to produce different mRNA isoforms from the same gene. Alternative splicing is a prevalent phenomenon that greatly expands proteome diversity and that also contributes to the quantitative regulation of gene expression (3,4). The binding of RBPs, like serine/arginine-rich (SR) proteins and hnRNP proteins, to short degenerated RNA motifs located within exons or in their flanking introns, is determinant for the control of alternative exon inclusion (5,6). The activity of a given RBP can also be favoured or antagonized by other factors, hereafter defined generally as splicing factors (SF), and their joint action or competition can promote or inhibit the assembly of the spliceosome onto nearby splice sites (7,8).

The specificity and intrinsic properties of each SF, even those of general constituents of the spliceosome, combined to the unique sequence and structural organization of each exon, explain why the knockdown of a specific factor in cells only results in the differential splicing of a limited number or transcripts, highlighting the relative plasticity of the spliceosome. The identification and quantification of splicing changes controlled by a given factor can be monitored by RNA sequencing (RNA-seq) technologies (9). However, it is more difficult to understand or to predict the cooperative or antagonistic effects of SFs on alternative splicing, especially on a genome-wide scale.

To address this question, we present SplicingLore, a database and web resource that contains the lists of alternative exons that are differentially included upon knockdown of 75 different SFs in various human cell lines. This collection is integrated into a website from which the user can easily download a given dataset of SF-regulated exons or submit a single exon or a list of exons to predict their potential regulation by the SFs of the database. As a proof of concept of the utility of our resource, we predicted and experimentally validated a new functional cooperation between RNA helicases DDX17 and DDX5 and the HNRNPC protein. SplicingLore is freely available at https://splicinglore.ens-lyon.fr/.

## MATERIALS AND METHODS

### Database and web interface

SplicingLore database was implemented on Linux Ubuntu 22.04, using a MySQL database (www.mysql.com) and organized in 9 tables. The web interface was developed using the programming languages PHP, CSS and JavaScript. Tools performing statistical tests were driven in R (v4.2.0) using R packages Tidyverse (<u>doi.org/10.21105/joss.01686</u>) or in Python (v3.10), using Plotly (10) for graphics.

### Data collection

SplicingLore is a database of 160 public datasets retrieved from GEO omnibus (11) and ENCODE consortium (12), linked to a user-friendly web interface (Table S1). All datasets were derived from Illumina RNA-seq experiments performed in 21 different human cell lines (293T, 786-O, A498, A673, Endoc-BH1, GM19238, HeLa, HepG2, SH-SY5Y, HMLE, Huh-7, K562, MCF7, MDA-MB-231, MG63, RD, RH30, RH41, RKO, LNCaP, LM2) in which the expression of 75 splicing factors was inhibited by siRNA or shRNA treatment.

### Differential exon inclusion analysis

For each dataset, we performed a qualitative and quantitative analysis of splicing variations using the FaRLine tool (13) (hg19 genome), comparing the condition in which the expression of the splicing factor was inhibited to the control condition. The following parameters were used to define exons that were differentially included: |ΔPSI| ≥ 10%, adjusted *p*-Val-≤ 0.05. The ΔPSI corresponds to the change in exon inclusion (PSI: percent spliced-in) between two conditions. We then retrieved and fed the SplicingLore database with the lists of differentially included cassette exons corresponding to each dataset.

### Randomization test and scoring

#### - Input list of exons with ΔPSI and *p*-values

We defined a Pearson correlation between the ΔPSI of two sets of exons. The computed empirical *p*- value is the probability of observing a Pearson correlation as high or higher as the observed correlation when considering a set of randomly associated exons. This probability was computed from the empirical cumulative distribution function generated by computing the Pearson correlation for 10^4^ random sets of exon (which leads to a maximal *p*-value resolution of 10^−4^).

The “percent.sig.input” is the fraction of significantly regulated exons from the input list of exons that are also found in the list of exons regulated by the indicated SF. The “percent.sig.SF” is the fraction of significantly regulated exons from the list of exons regulated by the indicated SF that are also found in the input list of exons.

A confidence score (Score) was set up to facilitate the identification of candidate SF. It is linked to a permutation statistic test, which also gives a *p*-value, the “percent.sig.input” and the “percent.sig.SF”. The Score ranges between 0 and 1, which indicates the degree of correlation or anti-correlation between the effect of a given SF and the input list of exons provided by the user.

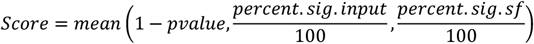

#### - Basic input list of exons (chromosome coordinates only)

For an input set of exons *E*, randomization tests were performed to test if the number of exons *N* regulated by a given SF *S* is enriched or impoverished. For this, 10^4^ sets of control exons with the same size as *E* were sampled. Then the number of exons regulated by *S* was computed for each control set. Finally, an empirical *p*-value was computed for S as:

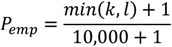

where k is the number of controls sets with a number of exons regulated by *S* higher or equal to *N*, and *l* is the number of controls sets with a number of exon regulated by S lower or equal to *N*. This *p*-value was computed for each SF and was then corrected using the Benjamini-Hochberg procedure.

### Cell culture and transfections

Human embryonic kidney 293T cells were grown as recommended by the manufacturer and transfected as described previously (14). For knockdown experiments, we used a total of 40 nM siRNA as follows : 40 nM siCtrl for control experiments, 20 nM of si*DDX5/DDX17* or si*HNRNPC* + 20 nM siCtrl for single factor depletion, or 20 nM si*DDX5/DDX17* + 20 nM si*HNRNPC* for double knockdown. Cells were harvested 48h later. Sequences of siRNAs are given in Table S2.

### Co-immunoprecipitation and Western blotting

Total protein extraction was carried out as previously described (15). Primary antibodies used for Western-blotting: DDX5 (ab10261, Abcam), DDX17 (ab24601, Abcam), HNRNPC (D6S3N, Cell Signaling), GAPDH (sc-1616, SantaCruz).

For co-immunoprecipitation, cells were harvested and gently lysed for 5 minutes on ice in a buffer containing 10 mM Tris-HCl pH 8.0, 140 mM NaCl, 1.5 mM MgCL_2_, 10 mM EDTA, 0.5% NP40, completed with protease and phosphatase inhibitors (Roche #11697498001 and #5892970001), to isolate the nuclei from the cytoplasm. After centrifugation, the nuclei were lysed in the IP buffer (20 mM Tris-HCl pH 7.5, 150 mM NaCl, 2 mM EDTA, 1% NP40, 10% Glycerol and protease/phosphatase inhibitors) for 30 minutes at 4°C under constant mixing. The nuclear lysate was centrifuged for 15 minutes to remove debris and soluble proteins were quantified by BCA (Thermo Fisher Scientific). The lysate was pre-cleared with 30 μL of Dynabeads Protein G (Thermo Fisher Scientific) for 30 minutes under rotatory mixing, and then split in 1.5 mg aliquots of proteins for each assay. Each fraction received 5 µg of antibody and the incubation was left overnight at 4°C under rotation. The following antibodies were used for IP: rabbit anti-DDX17 (19910-1-AP, ProteinTech) and anti-HNRNPC (PA522280, Thermo Fisher Scientific) or a control rabbit IgG (Thermo Fisher Scientific), goat anti-DDX5 (ab10261, Abcam) or control goat IgG (Santa Cruz). The next day, the different lysate/antibody mixtures were incubated with 50 µl Dynabeads Protein G (Thermo Fisher Scientific) blocked with bovine serum albumin, for 4 hours at 4°C under rotation. Bead were then washed 5 times with IP buffer. Elution was performed by boiling for 5 minutes in SDS-PAGE loading buffer prior to analysis by Western-blotting.

### RNA extraction and PCR analyses

Total RNA were isolated using TriPure Isolation Reagent (Roche). For reverse transcription, 2 µg of purified RNAs were treated with Dnase I (Thermo Fisher Scientific) and retrotranscribed using Maxima reverse transcriptase (Thermo Fisher Scientific), as recommended by the manufacturer. Potential genomic DNA contamination was systematically verified by performing negative RT controls in absence of enzyme, and by including controls with water instead of cDNA in PCR assays. All PCR analyses were performed on 0.5 ng cDNA using 0.5 U GoTaq® DNA polymerase (Promega). Quantification of PCR products was performed using the Image Lab software (BioRad) after agarose gel electrophoresis. The PSI (percent spliced-in) value of each alternative exon was calculated in each condition using the following formula: inclusion product / (inclusion product + skipping product) x100. The ΔPSI corresponds to the difference between the PSI for each silencing condition and the PSI of the siCtrl condition. Sequences of all primers are given in Table S2.

## RESULTS

### Data collection and analysis

To feed the SplingLore database, we searched the GEO omnibus (10) and ENCODE (11) databases for datasets generated from Illumina RNA seq experiments that all consisted in knocking down a given splicing factor (SF) in a human cell line, by means of a treament with siRNA or shRNA (Figure 1A). We filtered out the datasets for which the sequencing quality or depth was too low to reach our objective, which was to obtain robust and reliable lists of alternative exons regulated by the tested SF. We eventually obtained a curated list of 160 datasets, corresponding to 75 SFs tested in 21 different cell lines (Figure 1A, Table S1). Three cell lines (HepG2, K562 and 293T) represented more than 75% of the datasets (Figure 1B), and some SFs were tested in up to 5 cell lines.

**Figure 1.**
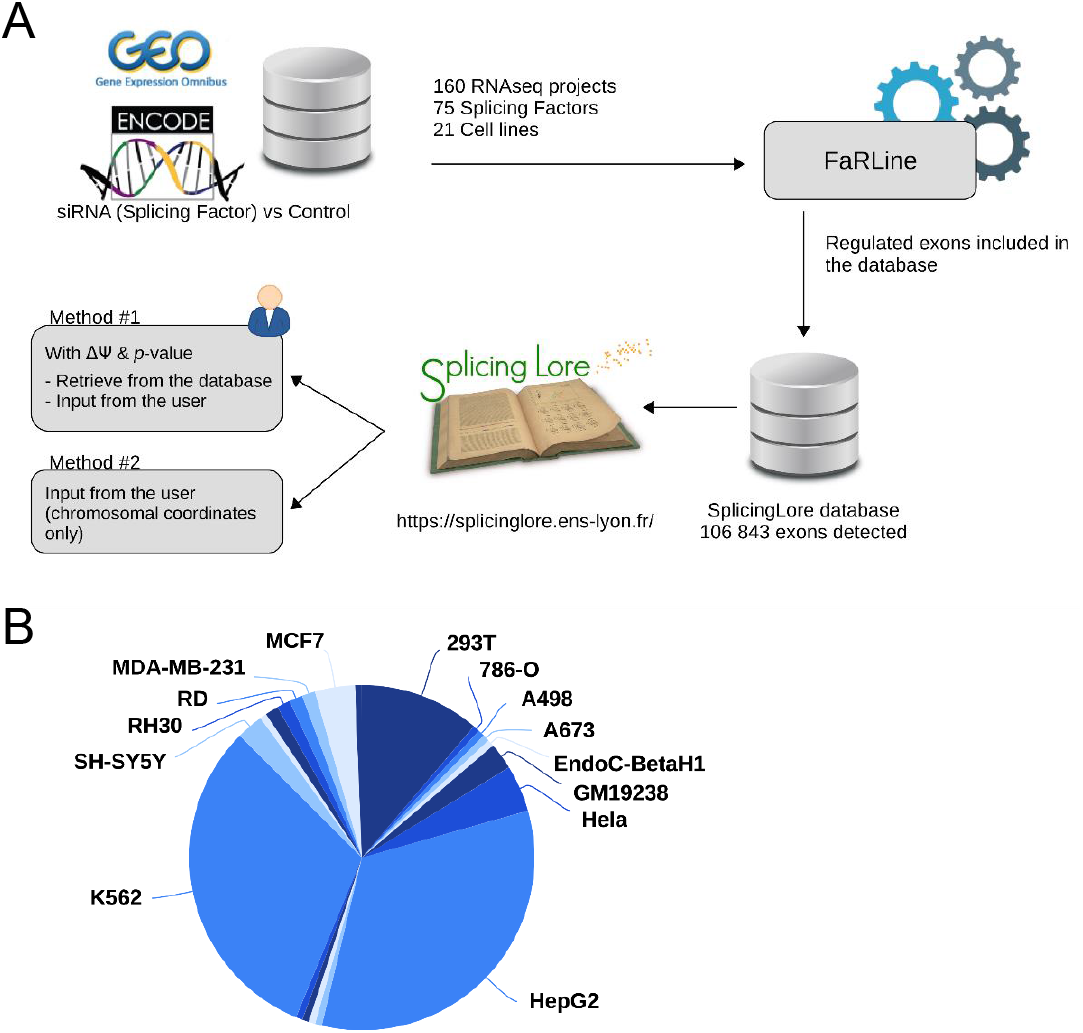
General features of SplicingLore. **A**. Outline of the SplicingLore workflow. RNA-seq datasets from GEO and ENCODE databases were uniformly processed by the FaRLine tool to generate lists of alternative exons regulated by 75 splicing factors in 21 human cell lines. These datasets can be used to compute correlation scores of exon regulation with a list of exons of interest, which is either retrieved from the database or provided by the user (query Method 1). The user can alternatively query SplicingLore with a list of exons associated only with their chromosomal coordinates (Method 2). In the latter case, the list of exons can include or not a ΔPSI and *p*-value. **B**. Diagram showing the repartition between cell lines of the datasets stored in SplicingLore.

The selected datasets were all processed in the same way using FaRLine, a splicing-dedicated pipeline developed earlier in our lab (13). We thus obtained uniformly formatted lists of alternative splicing events for each dataset. Although FaRLine detects and quantifies variations in alternative 5’ and 3’ splice sites, as well as multiply spliced exons, we decided to focus only on single cassette exons, which represent the vast majority of regulated splicing events and which are easier to handle when comparing multiple datasets. Note that the annotation of all exons is based on the FasterDB database (https://fasterdb.ens-lyon.fr/faster/home.pl) (16).

### Web interface and graphic visualisation

SplicingLore can be queried in two different ways (Figures 1A and 2A). The first method allows the user to analyse a list of exons associated with parameters of splicing regulation. The default format requested for this analysis includes the gene symbol and exon number, chromosomal coordinates, as well as values of differential exon inclusion (ΔPSI) and *p*-value (Method 1, Figure 2A, left window). Gene symbols and exon numbers correspond to gene annotations of the FasterDB database (16), but the critical details to provide here are the chromosomal coordinates. Incorrect gene symbol and/or exon number will not impede the search but instead will issue a warning message and a proposition for correcting the input, based on coordinates (see below).

**Figure 2.**
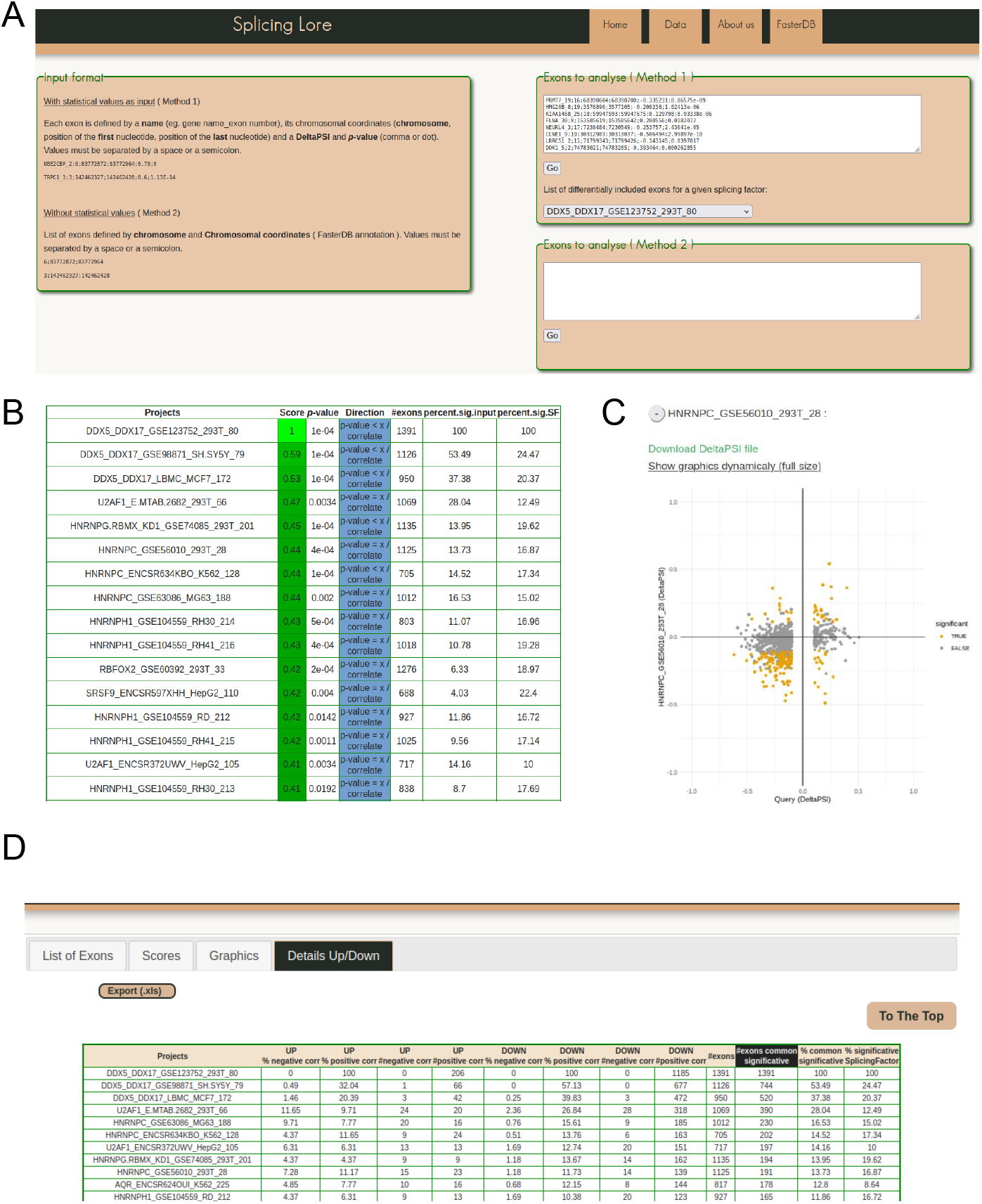
Visualization of the SplicingLore web Interface. **A**. View of the home page, with the instructions to prepare the input list of exons and the query window. This includes also the drop-down menu that allows to retrieve a given list of SF-regulated exons from the database. **B**. View of the top of the main output file, showing for each dataset the different correlation parameters with the query list (here the exons regulated by DDX5/DDX17 in 293T cells). The Score and associated *p*-value for positive or negative correlation are provided for each dataset of the database, along with the total number of exons in the corresponding dataset and the fraction of common exons between the compared datasets, relative to the query list or to the tested dataset. **C**. Correlation graph of the ΔPSI values for common exons from the query list and one of the stored datasets. Orange dots represent exons that are significantly regulated by both SF. The file containing the corresponding ΔPSI values for these exons can be downloaded here. **D**. View of the top of the detailed output file, which provides the number (#) and fraction (%) of shared exons between the query and the dataset, separated in up-regulated and down-regulated sub-classes of exons.

The query list of exons can be automatically retrieved from one of the projects referenced in the database, upon selection of the dataset of interest from the drop-down menu (Figure 2A, top right window). Alternatively, the user can upload a list of exons of interest to query SplicingLore for their potential regulation by the SFs of the database. As each exon is associated to a ΔPSI value, this first method will allow to look for positive and negative correlations between the input regulation and the regulation by SFs from the database.

The second query method is less restrictive as it does not require any parameter of exon inclusion (ΔPSI and *p*-value), if these values are not available (Method 2, Figure 2A). In this mode, the user can upload a list of exons that are only identified by their chromosomal coordinates (Figure 2A, bottom right window).

Once the query is launched, the first displayed page (“List of exons” tab) informs the user about exons which were not properly recognized, either because they were not detected in any of the 160 datasets, or because they were incorrectly formatted. In the latter case, corrected features are given to allow the user to restart the analysis (Figure S1).

From the “Scores” tab, the user obtains a downloadable table (Figure 2B) which recapitulates the correlation scores between the query list and each of the SplicingLore datasets, as well as different comparison parameters that are explained below. Since this table is sorted based on our custom Score, the user can immediately visualize on top of the list which SF is more susceptible to be involved in the regulation of the tested exons. Scores indicating a significant positive or negative correlation are shown in different colours to facilitate their rapid visualization on the website (downloaded files are colourless). The “percent.common.query” and “percent.common.SF” columns indicate the fraction of common exons between the compared datasets, calculated from the query list or from the tested dataset, respectively (see Figure S2 for a representation of these fractions).

The “Graphics” tab presents the same information in the form of a correlation graph, which is available in a fixed or interactive mode (Figure 2C). The values used to generate these graphs can also be downloaded as a table. Finally, a more comprehensive table is available in the “Details Up/Down” tab (Figure 2D). Alongside the information described above, it also provides the number (#) and fraction (%) of exons that are shared by both compared datasets, taking into account the direction of their regulation (up-regulation or down-regulation of their inclusion upon SF knock-down).

### Example of application

In order to check out the validity of our tool, we retrieved from SplicingLore the list of exons whose inclusion was altered upon the knock-down of DEAD box RNA helicases DDX5 and DDX17 in the 293T cell line (GSE123752, Table S3). We then used this list as a query to search for possible correlative (or anti-correlative) effects between these splicing regulators and other SFs.

The first lines of the resulting table show that beside the query list itself (score of 1), the best correlation scores correspond to the other two DDX5/DDX17 datasets included in SplicingLore, in SH-SY5Y and MCF7 cell lines, with respective scores of 0.59 and 0.53 (Figure 2B, Table S4). Interestingly, we also observed a good correlation (with a score of 0.41-0.43) between the tested DDX5/DDX17 dataset and five different HNRNPH1 datasets (Figure 2B, Table S4). This is in agreement with the fact that HNRNPH1 was previously described as a co-regulator of DDX5/DDX17-mediated splicing in myoblasts and epithelial cells (15).

Results from Table S4 also revealed HNRNPC as one of the top predicted factors, with positive correlation scores in 4 datasets (score= 0.44 in 293T, K562 and MG63 cell lines, and 0.38 in HepG2 cells) (Figure 2B, Table S4). When considering separately the subclasses of up- and down-regulated exons, we noticed that the positive correlation between DDX5/DDX17 and HNRNPC exons was especially visible for down-regulated exons (Table S5, Figure S3). Indeed, up to 15.6% of exons down-regulated upon HNRNPC knockdown were regulated in the same manner (11.5% in average for the 4 correlated datasets), while only about 1% were regulated in the opposite manner (Tables S5 and S6). For up-regulated exons, results were more variable between the different datasets, although a trend for a positive correlation was also observed, especially in 293T and K562 cells. These 2 datasets displayed more than 11% of positive correlation with the query, these values being the highest among all datasets (apart from DDX5/DDX17 datasets, Table S5).

To explore further the possibility of a functional link between DDX5/DDX17 and HNRNPC, we first queried SplicingLore in a reverse manner, uploading on the website a list of 3030 exons corresponding to the union of all exons regulated upon HNRNPC knock-down in 4 datasets of our database (293T, K562, HepG2 and MG63). This analysis identified the 3 DDX5/DDX17 datasets among those showing a significant positive correlation (scores from 0.46 to 0.50) with the query list (Table S7, Figure S4).

Finally, to experimentally validate our predictions, we silenced the expression of DDX5/DDX17 and HNRNPC in 293T cells, independently or together (Figure 3A and Figure S5A), and we monitored the effect of these treatments on a selection of alternative exons whose inclusion was found to be impacted by the knock-down of these factors (10 more skipped exons and 7 more included exons). In conditions of single depletion, the inclusion of all exons was modified according to the predictions and interestingly, the combined depletion of DDX5/DDX17 and HNRNPC significantly enhanced the effect of single siRNA treatment on splicing (Figures 3B and 3C). This suggested that DDX5/DDX17 and HNRNPC could cooperate to regulate exon inclusion, which prompted us to test whether these factors interact with each other. Indeed, endogenous HNRNPC and DDX17 co-immunoprecipitated in an RNA-independent manner, suggesting a possible direct interaction (Figure 3D). No clear sign of interaction was observed between DDX5 and HNRNPC (Figure S5B).

**Figure 3.**
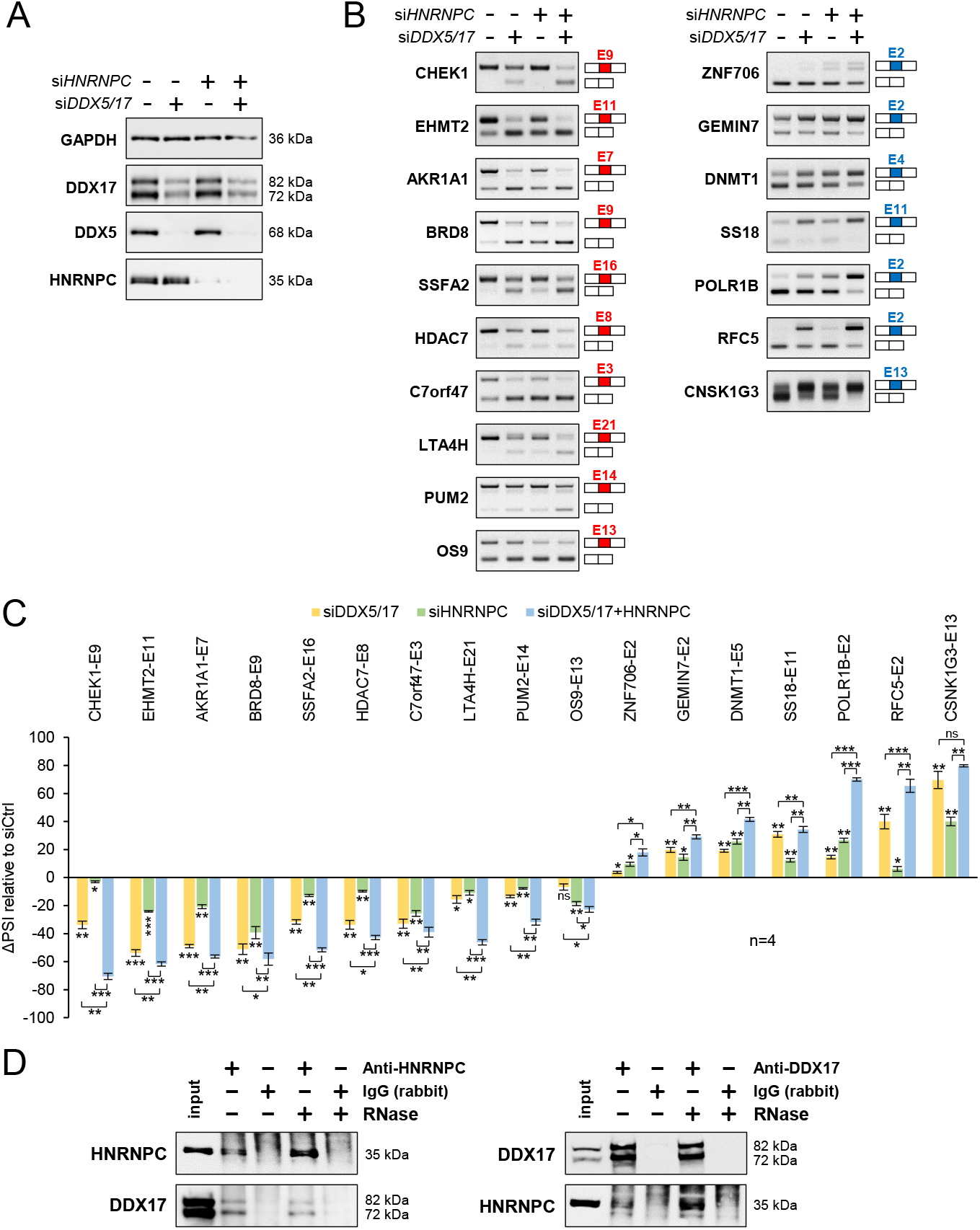
Validation of a functional relationship between DDX5/DDX17 and HNRNPC. **A**. Western-blot showing the expression of DDX5, DDX17 and HNRNPC following treatment with siRNA targeted against luciferase (negative control), DDX5/DDX17 and HNRNPC. Quantification of this experiment is shown in Figure S5A. **B**. RT-PCR analysis showing the inclusion of a selection of alternative exons after the depletion of DDX5/DDX17 and/or HNRNPC in 293T cells. The corresponding gene and exon number (according to FasterDB annotation) is indicated. Exons down-regulated and up-regulated upon SF knock-down are represented in red and blue, respectively. **C**. Quantification of the RT-PCR experiment shown in C. The indicated ΔPSI values correspond to the difference between the PSI (percent spliced-in) value of each depleted sample and the control sample. Statistical comparison between each condition (including the unshown control condition) was calculated using a one-way ANOVA (Holm–Sidak’s multiple comparison tests: * P-val < 0.05; ** P-val < 0.01; *** P-val < 0.001). **D**. Co-immunoprecipitation assays between endogenous HNRNPC and DDX17 in 293T cells, in the absence of presence of RNase A.

Altogether, these results disclosed a novel functional partnership between HNRNPC and the helicase DDX17, and illustrated the usefulness of SplicingLore to improve our knowledge on splicing regulation.

## DISCUSSION

One challenge faced by scientists from the alternative splicing field is to decode the cooperative or antagonistic effects of SFs to understand and eventually predict splicing outcomes on a genome-wide scale (17). Exploiting a large collection of RNA-seq-derived lists of exons regulated by 75 SFs, SplicingLore helps to fill this knowledge gap in a straightforward manner. This new resource enables cross-comparisons between datasets, or allows to search for SFs that control the inclusion of a list of exons provided by the user. This resource only requires to format correctly the list of exons of interest, and it is therefore fast and easy to run. All datasets were selected based on the quality and coverage depth of the sequencing experiment, and they were processed in a standardized manner. This unique experimental workflow ensures a reliable calculation of correlation scores between the tested dataset and good validation efficiency when testing the inclusion of the predicted exons experimentally.

As a proof of concept, we searched for SFs that could stimulate or antagonize the inclusion of alternative exons regulated by RNA helicases DDX17 and DDX5. These closely related paralog proteins belong to the large family of evolutionarily conserved DEAD-box ATP-dependent RNA helicases (18). Previous reports, from our lab and others, have shown that DDX5 and DDX17 control the inclusion of a large number of exons by modulating the folding of their target transcripts, thanks to their helicase activity, and by modulating the binding of splicing regulators to RNA (14,15,19-23). This is the case of HNRNPH for example, whose binding to RNA is facilitated by DDX5/DDX17 (15), and which was predicted by SplicingLore among the top predicted SFs for positive correlation with the helicases.

SplicingLore predicted a good correlation in the regulation of several hundreds of exons by DDX5/DDX17 and HNRNPC, including exons that are either activated or repressed by those factors. We experimentally validated the additive effect of these factors on a subset of both classes of exons, and found that HNRNPC and DDX17 associate with each other in cells in an RNA-independent manner. The molecular nature of this functional relationship is not clear, and understanding how HNRNPC stimulates exon inclusion while it is often described as a splicing repressor (6) will require further investigation.

In conclusion, SplicingLore represents a quick and easy tool to investigate alternative splicing regulation by a large panel of SFs. Beside the specialists of this field, the straightforwardness of this new resource will also make it more widely appreciated by scientists who are less familiar with the complex interplays that underlie splicing regulation.

## AUTHOR CONTRIBUTION

Conceptualization: H.P., D.A. and C.F.B.; methodology and statistical analysis: H.P and N.F.; data collection and processing: H.P., A.L. and N.F.; experimental validation and interpretation: J.V. and C.F.B.; software: H.P., N.F. and S.J.; writing of the original draft: H.P. and C.F.B.; review & editing: H.P., F.M., D.A. and C.F.B.

## ACKNOWLEDGEMENTS

I-Stem is part of the Biotherapies Institute for Rare Diseases (BIRD) supported by the Association Française contre les Myopathies (AFM-Téléthon). We gratefully acknowledge support from the PSMN (Pôle Scientifique de Modélisation Numérique) of the ENS of Lyon for computing resources. We also thank Laurent Modolo and other members of the LBMC Biocomputing Hub for helpful discussions.

## FIGURE LEGENDS

**Figure S1.**
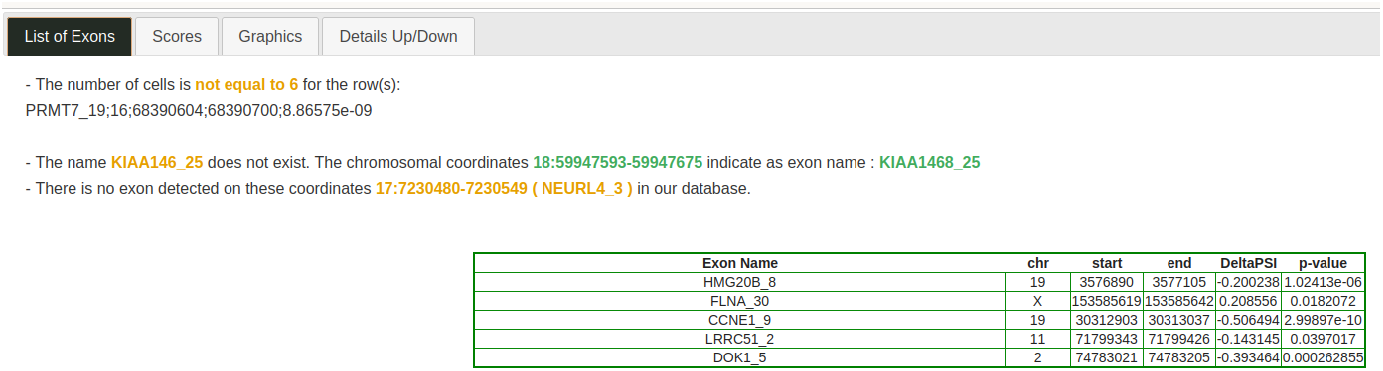

**Figure S2.**
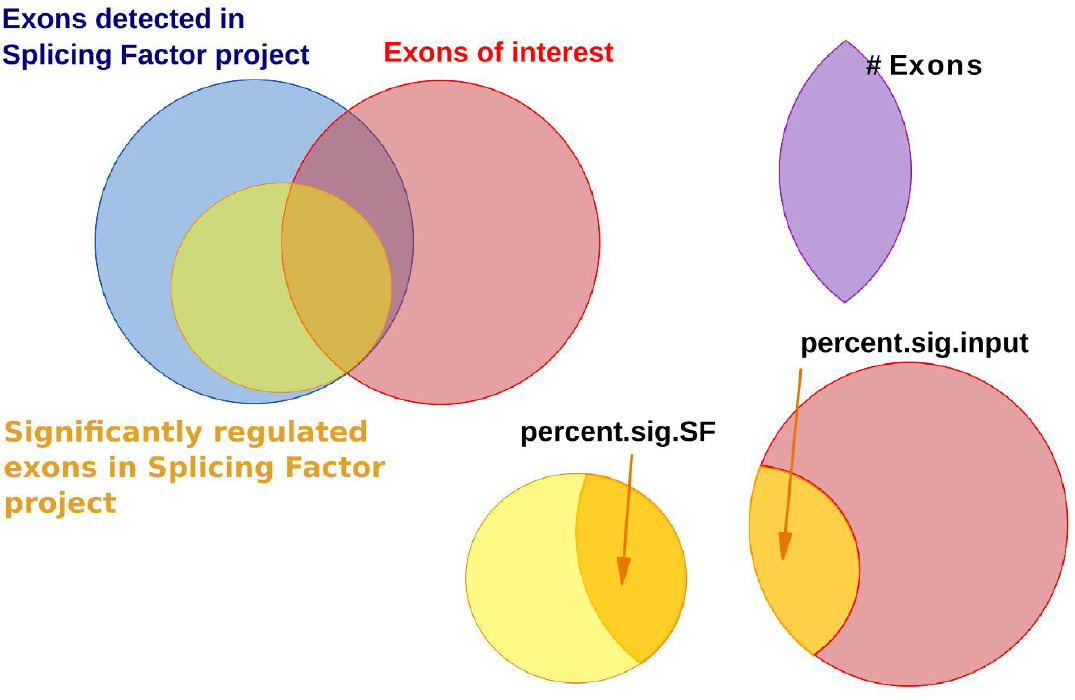

**Figure S3.**
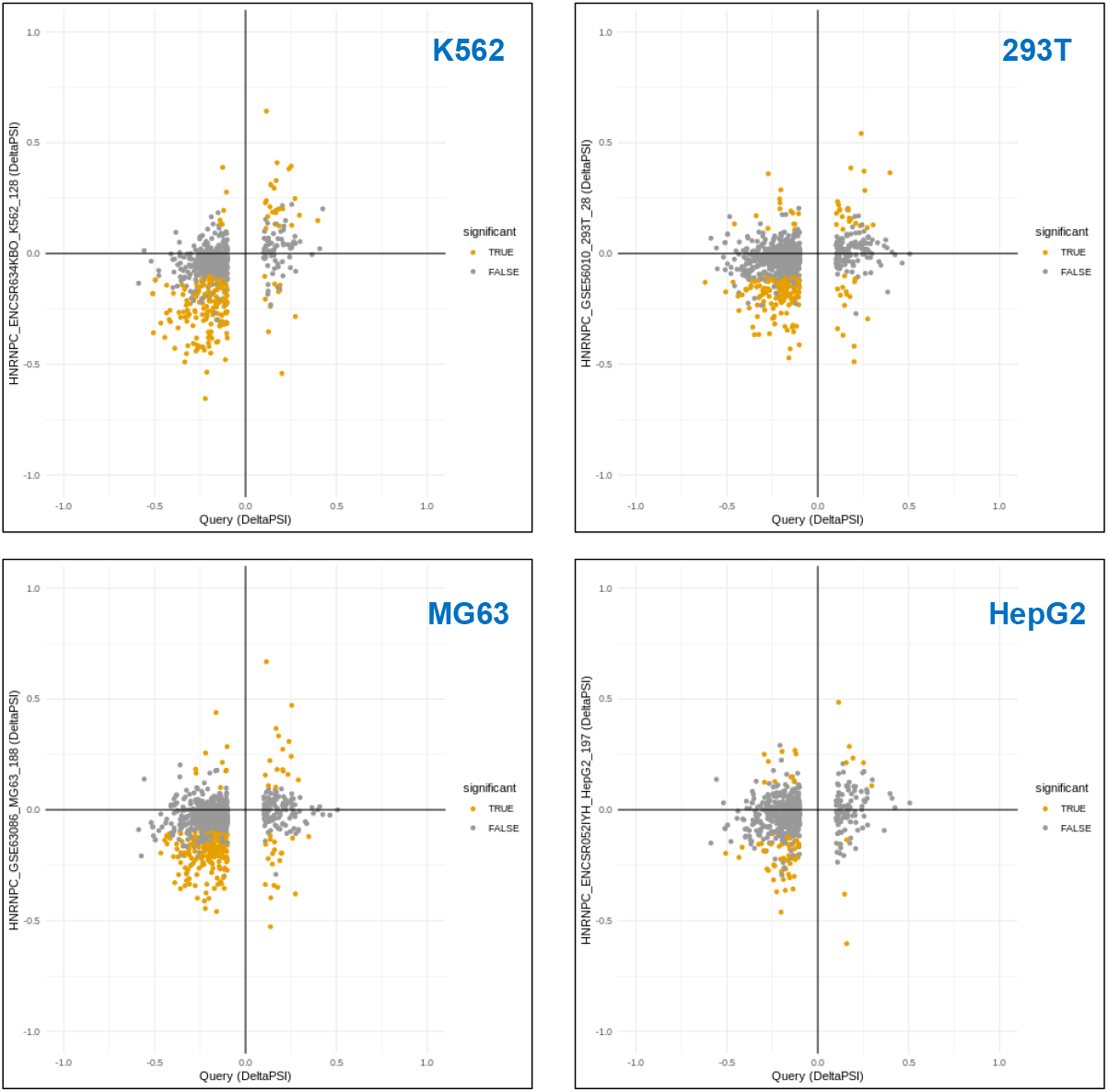
Correlation graphs between the query (DDX5/DDX17_293T) and the different HNRNPC datasets.

**Figure S4.**
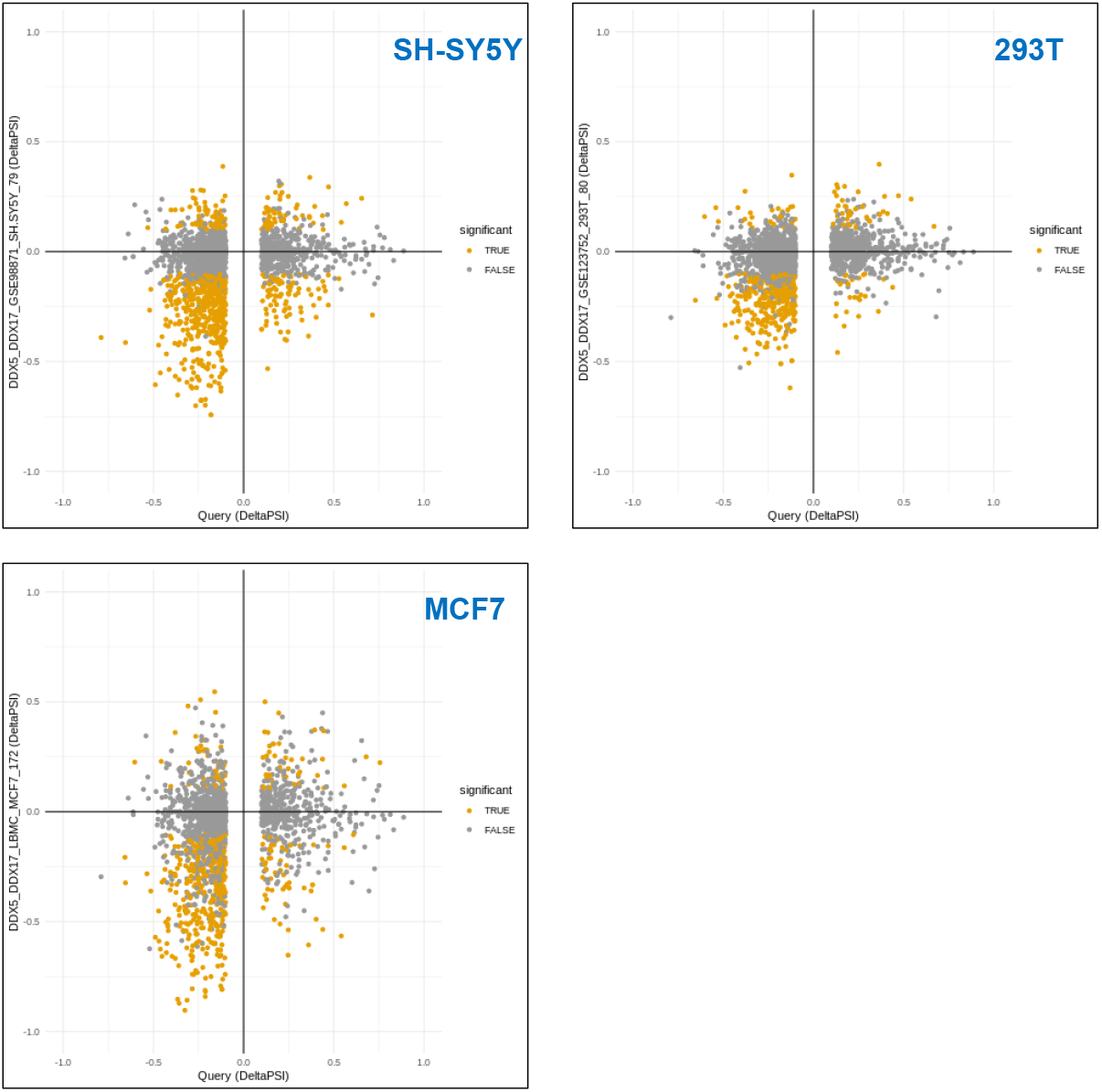
Correlation graphs between the query (HNRNPC_union) and the different DDX5/DDX17 datasets.

**Figure S5.**
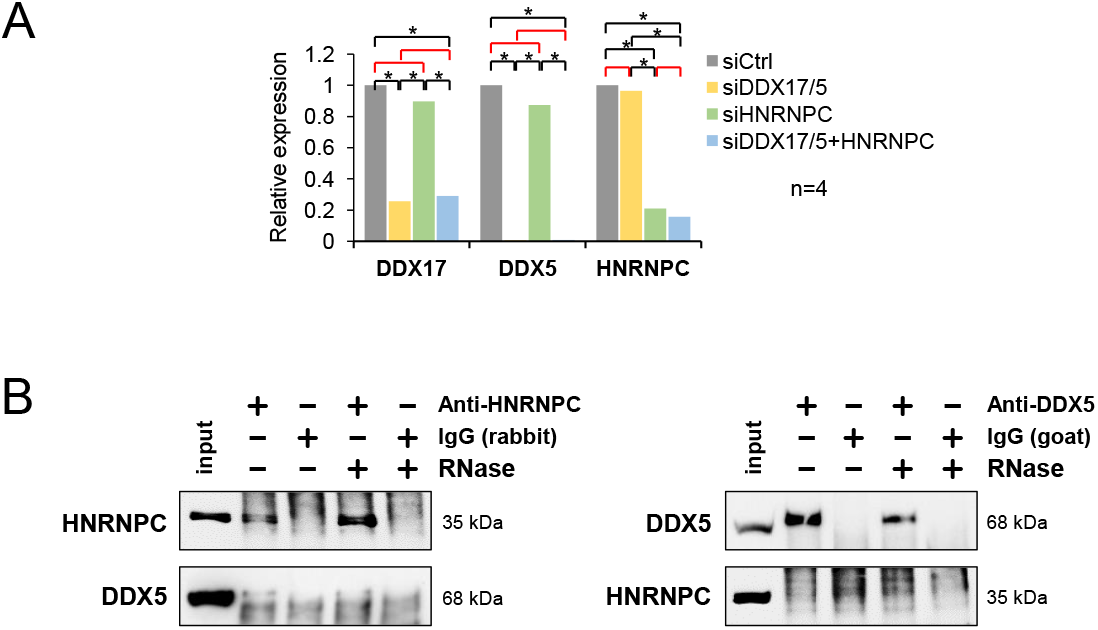
**A.** Quantification of the Western-blot experiment shown in Figure 3B. Measured signals for each protein were normalized to the GAPDH level (mean value of 4 independent experiments ± S.E.M.). Statistical test: one-way ANOVA (Holm–Sidak’s multiple comparison tests: * P-val < 0.01; non-significant comparisons are marked in red). **B**. Co-immunoprecipitation assays between endogenous HNRNPC and DDX5 in 293T cells, in the absence of presence of RNase A.

